# Occasional and constant exposure to dietary ethanol shortens the lifespan of worker honey bees

**DOI:** 10.1101/2024.04.03.586924

**Authors:** Monika Ostap-Chec, Daniel Bajorek, Weronika Antoł, Daniel Stec, Krzysztof Miler

## Abstract

Honey bees (*Apis mellifera*) are one of the most crucial pollinators, providing vital ecosystem services. Their development and functioning depend on essential nutrients and substances found in the environment. While collecting nectar as a vital carbohydrate source, bees routinely encounter low doses of ethanol from yeast fermentation. Yet, the effects of repeated ethanol exposure on bees’ survival and physiology remain poorly understood. Here, we investigate the impacts of constant and occasional consumption of food spiked with 1% ethanol on honey bee mortality and alcohol dehydrogenase (ADH) activity. This ethanol concentration might be tentatively judged close to that in natural conditions. We conducted an experiment in which bees were exposed to three types of long-term diets: constant sugar solution (control group that simulated conditions of no access to ethanol), sugar solution spiked with ethanol every third day (that simulated occasional, infrequent exposure to ethanol) and daily ethanol consumption (simulating constant, routine exposure to ethanol). The results revealed that both constant and occasional ethanol consumption increased the mortality of bees, but only after several days. These mortality rates rose with the frequency of ethanol intake. The ADH activity remained similar in bees from all groups. Our findings indicate that exposure of bees to ethanol carries harmful effects that accumulate over time. Further research is needed to pinpoint the exact ethanol doses ingested with food and exposure frequency in bees in natural conditions.

## Introduction

Insect pollinators play a crucial role in the functioning of both natural and agricultural ecosystems (Klein et al. 2007, Gallai et al. 2009). During pollination services, they collect pollen, essential for growth, and nectar, a primary energy source. The quality and composition of their diet are critical for development, health, and survival, making the abundance and diversity of resources in the surrounding environment crucial. While collecting pollen and nectar, pollinators are exposed to substances that directly impact them. For instance, insecticides widespread in agricultural environments significantly increase mortality and adversely affect the development and foraging abilities of pollinators (e.g. in solitary bees, Boff et al. 2021, Mokkapati et al. 2021). Conversely, certain phytochemicals found in pollen and nectar, such as caffeine, can significantly reduce parasitisation, acting both preventatively and therapeutically on pollinators’ health (e.g. in bumble and honey bees, Folly et al. 2021, Motta et al. 2023). Pollinators can be passively affected by environmental substances due to ingestion of contaminated resources, but they can also actively seek dietary components. For example, honey bees actively select food types advantageous to them in specific situations, such as opting for nectar with higher antibiotic potential during infection (Gherman et al. 2014) or gathering propolis to combat rising parasite levels (Pusceddu et al. 2021).

Honey bees (*Apis mellifera* L.), which are one of the key insect pollinators, likely encounter and consume ethanol in their natural environment. The main carbohydrate source for honey bees, floral nectar, is frequently colonised by yeasts capable of producing ethanol through fermentation (Lievens et al. 2015). Observations from tropical regions indicate that the concentration of ethanol in palm flower nectar can reach 6.9% (Hockings et al. 2015, Gochman et al. 2016). A concentration of about 1% ethanol, especially in more temperate climates, seems more realistic as a component of nectar in natural conditions (Ehlers and Olesen 1997, Jakubska et al. 2005, Goodrich et al. 2006, Wiens et al. 2008, Golonka et al. 2014, Rering et al. 2018). In any case, ethanol consumption must be significant enough to allow honey bees to biosynthesise ethyl oleate — a crucial pheromone that regulates colony demography by inhibiting the transition of workers into foragers (Leoncini et al. 2004, Castillo et al. 2012a, Castillo et al. 2012b). Only foragers, the type of individuals that leave the hive and explore the surroundings for food, possess dehydrogenase enzymes likely responsible for breaking down ethanol (Miler et al. 2021). Furthermore, despite the aversive taste of ethanol in water for honey bees, when dissolved in sugar, it is willingly consumed both in the laboratory and field, even in extreme concentrations as high as 20% (Abramson et al. 2000, Abramson et al. 2004a, Maze et al. 2006, Sokolowski et al. 2012, Mustard et al. 2018). Moreover, foragers prefer feeding solutions containing about 1% or even 2.5% ethanol over pure sugar solutions (Mustard et al. 2018, Varnon et al. 2018). All this suggests not only that ethanol frequently contaminates carbohydrate resources utilised by honey bees but also that they need this dietary component and are adapted for its encounter.

Nevertheless, ethanol has been observed to induce multiple negative effects on honey bees, both in social and non-social contexts. It disrupts social communication among individuals by directly influencing behaviours such as antennation or trophallaxis (Mixson et al. 2010, Wright et al. 2012) and through increasing aggression (Abramson et al. 2004b, Ammons and Hunt 2008). Moreover, ethanol consumption in foragers modifies proper dance communication (Bozic et al. 2006). Ethanol also impairs honey bees’ locomotion, foraging, and learning (Abramson et al. 2005, Giannoni-Guzmán et al. 2014, Maze et al. 2006, Mustard et al. 2008, Black et al. 2021, Ahmed et al. 2022). The extent of these behavioural changes occurs in a dose-dependent manner, with higher doses of ethanol producing larger effects on behaviour (Maze et al. 2006, Bozic et al. 2007, Wright et al. 2012). However, the majority of the studies that document effects resulting from ethanol consumption focus on single-exposure impact, often with concentrations that are relatively high and unlikely of ecological relevance. Not only the effects of low ethanol levels are understudied but so are those related to repeated ethanol exposure. Thus far, it has been demonstrated that bees show tolerance, expressed as lower motor impairment in response to ethanol in individuals previously exposed to it than those exposed for the first time (Miler et al. 2018 but see Stephenson et al. 2021). Moreover, individuals fed on food spiked with ethanol for a prolonged time exhibit withdrawal symptoms upon discontinuation of access to such food (Ostap-Chec et al. 2021). These characteristics can be considered primary hallmarks of alcohol dependence development in conditions of repeated encounters with ethanol. Still very little is known about the survivorship and physiological effects associated with repeated exposure to relatively low concentrations of ethanol in the diet.

Most research focuses on the behavioural and social effects of ethanol consumption in bees. However, these reactions are typically closely linked to physiological or biochemical responses. For instance, studies on *Drosophila* have shown that sensitivity and tolerance to ethanol are directly related to the activity of enzymes, such as alcohol dehydrogenase (ADH) (Ogueta et al. 2010). Research on vertebrates (Buris et al. 1985, Kishimoto et al. 1995, Tran et al. 2015) indicates that this enzyme tends to increase in activity with continuous, prolonged exposure to ethanol. In honey bee foragers, the presence of the most common type of ADH -type 1 - has been identified (Bouga et al. 2005, Miler et al. 2021, Miler et al. 2022). However, it is unclear whether the consumption of ethanol by foragers leads to increases in the activity of this enzyme.

Here, we investigate the effects of occasional and constant low-level ethanol consumption on the survival and ADH activity of forager-age honey bees. We hypothesise that there would be no significant increase in mortality compared to abstinent individuals. Moreover, we hypothesise that the ADH activity would be highest in individuals exposed to ethanol most frequently, lower in those with occasional exposure, and lowest in workers with no access to ethanol. Our expectations are based on current knowledge regarding the likely routine presence of ethanol in food resources utilised by bees, their preferences for this dietary component, and adaptations towards its encounter.

## Methods

### General procedure

We performed the experiment in 4 bouts, with 2 days of delay between each bout. For each bout, we obtained newly emerged individuals that originated from 2 different colonies. For that purpose, we took a single bee-free frame with a capped brood from each colony participating in a bout and placed it in an incubator (KB53, Binder, Germany) at 32 °C overnight. The next morning we marked emerged workers with a coloured dot on the thorax using a non-toxic paint marker and then released them into an unrelated hive. Leaving bees in the hive for a period after emergence allows for their proper development and immune system strengthening, resulting in high subsequent survival rates in laboratory cages. This enables conducting extended experiments, as demonstrated in a previous study by Ostap-Chec et al. (2021). When bees were 7 days old, we recollected marked workers by opening the hive and picking them frame-by-frame using forceps, placing them in wooden cages and transporting them to the laboratory. The workers were kept 100 individuals per cage, 9 cages per bout, and with access to *ad libitum* water and food (2 M sucrose solution) in an incubator (KB400, Binder, Germany) at 32 °C and 60-70% RH. We applied a period of acclimation that lasted for 2 weeks, to be sure that all workers reached forager age (Dukas 2008) and remained naïve in terms of ethanol presence in their previous diet. We noted exactly how many individuals were in each cage as there was some mortality during the acclimation period (the number of workers ranged from 94-100 by the point of reaching 21 days of age). Then, we created 3 replicates in each bout, with each replicate comprising 3 groups, and started the diet. For the control group (CONTROL), we provided the workers with a 2 M sucrose water solution. For the group with occasional exposure (OCCASIONAL), we provided the workers with the same solution, but every 3rd day, we switched it to a sucrose water solution with the addition of 1% ethanol. For the group with constant exposure (CONSTANT), we provided the workers with a sucrose water solution with the addition of 1% ethanol (Fig 1). Hence, we had 12 cages per group (4 bouts and 3 group replicates in each) totalling about 1200 individuals per group. The food with the addition of ethanol was prepared iso-caloric relative to the 2 M sucrose solution to avoid the confounding effect of ethanol caloric value when digested. Hence, assuming 5.5 kcal per 1 mL of ethanol, we added 1.38 g less sucrose to the food with ethanol to balance out the calories. Throughout the entire study, we refrained from providing protein sources to the workers and switched water and food to fresh every morning. During water and food renewal each day, we noted the number of dead individuals in each cage and removed the dead bodies. We exposed workers in each group to their respective diets for 21 days, and then the experiment was terminated. On the last day of the experiment, we froze 6 individuals from a randomly selected group replicate in each bout to analyse the alcohol dehydrogenase (ADH) activity in their bodies. Hence, we analysed 24 individuals per group in total. For that purpose, each body was thoroughly homogenised in 400 μl of PBS in a Bead Ruptor Elite homogenizer with ceramic beads (Omni International, USA) and spun in a centrifuge at 13000 RPM for 7 min. Then, 10 µl of each supernatant was analysed using the ADH assay kit (ab102533, abcam, the Netherlands) as per the manufacturer’s instructions. The kit enables the detection of type I ADH, which is present in bees (Miler et al. 2021). The optical density of the reaction product was read at 450 nm using the Multiskan FC microplate reader (Thermo Scientific, USA) after the initial incubation period (3 min) and in three measurement points between 7 and 35 min after the first measurement. The timeframe was determined to be within a range of linear reaction phase. The ADH activity (defined as nM NADH produced per min in 1 mL = mU/mL) was calculated based on the slope of the absorbance curve and translated into concentration values based on the prepared calibration curve slope. One sample was excluded from the analyses as the calculated activity had a slightly negative value (−0.087, 95% CI: -0.101-0.099), which was not biologically relevant.

**Figure 1.**
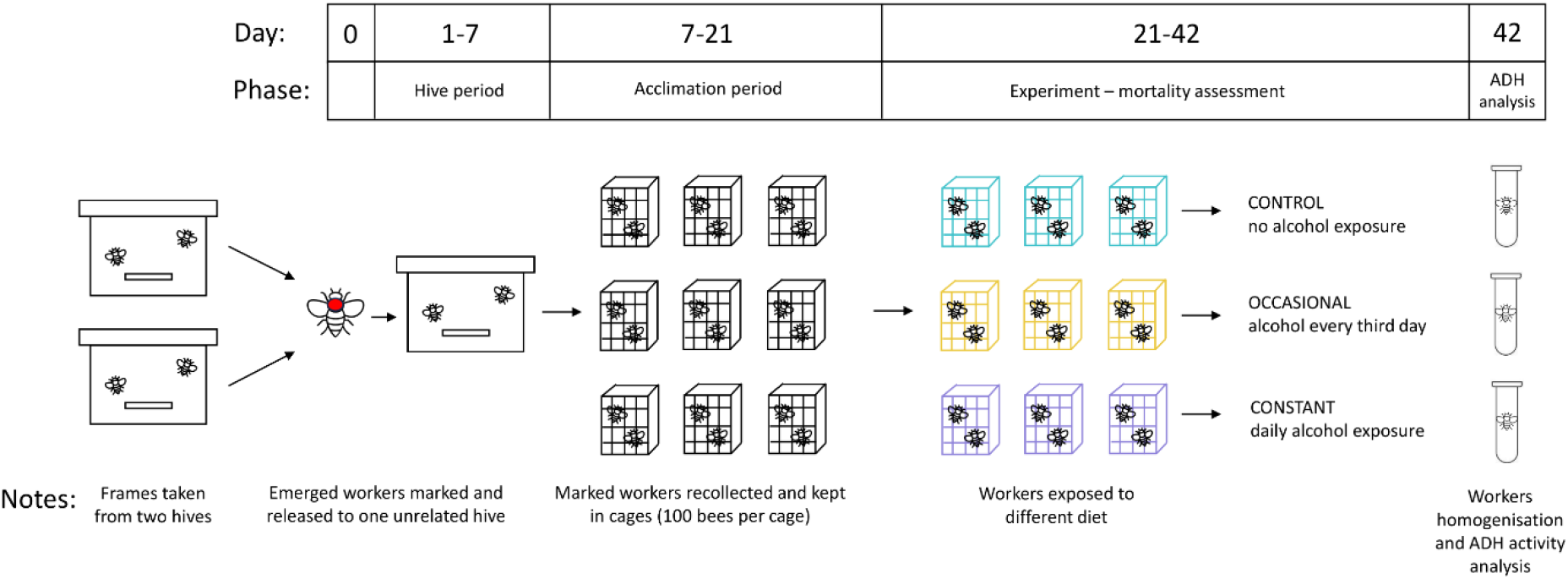
Experimental scheme. For clarity, the scheme presents the procedure for a single bout of bees (a mix of individuals from 2 colonies). We performed 4 bouts in total throughout the study.

### Statistics

To analyse the data, we used R (R Core Team 2024). For the analysis of survivorship, we used the Cox mixed-effect regression (‘coxme’ and ‘survival’ packages, Therneau 2022, 2023) with a fixed factor of the group and random factors of the bout and replicate (coxme function). Individuals that remained alive when the experiment was terminated (day 21 of the diet) were censored. To analyse the ADH activity, we used the generalised linear mixed-effects model (GLMM, ‘lme4’ package, Bates et al. 2015) with a Gamma distribution and a fixed factor of the group and a random factor of the bout (lme function).

## Results

Throughout the diet (21 days), the survival in the control groups (CONTROL) dropped to 79% (95% CI: 81-77%). Compared to that, the groups with occasional ethanol exposure (OCCASIONAL; Cox regression, z test = 7.24, p < 0.001) and with constant ethanol exposure (CONSTANT; Cox regression, z test = 18.02, p < 0.001) both demonstrated decreased survival. In the case of occasional exposure, the survival dropped to 65% (95% CI: 68-62%) whereas for constant exposure – to 41% (95% CI: 44-38%). Groups exposed to ethanol started to diverge from the control groups after 10 and 6 days of the diet for the OCCASIONAL and CONSTANT groups, respectively (Fig 2). The ADH activity did not differ between groups (GLMM: χ^2^ = 1.029, p = 0.598, Fig 3) and averaged 5.41 (SD: 3.16) mU/mL across groups.

**Figure 2.**
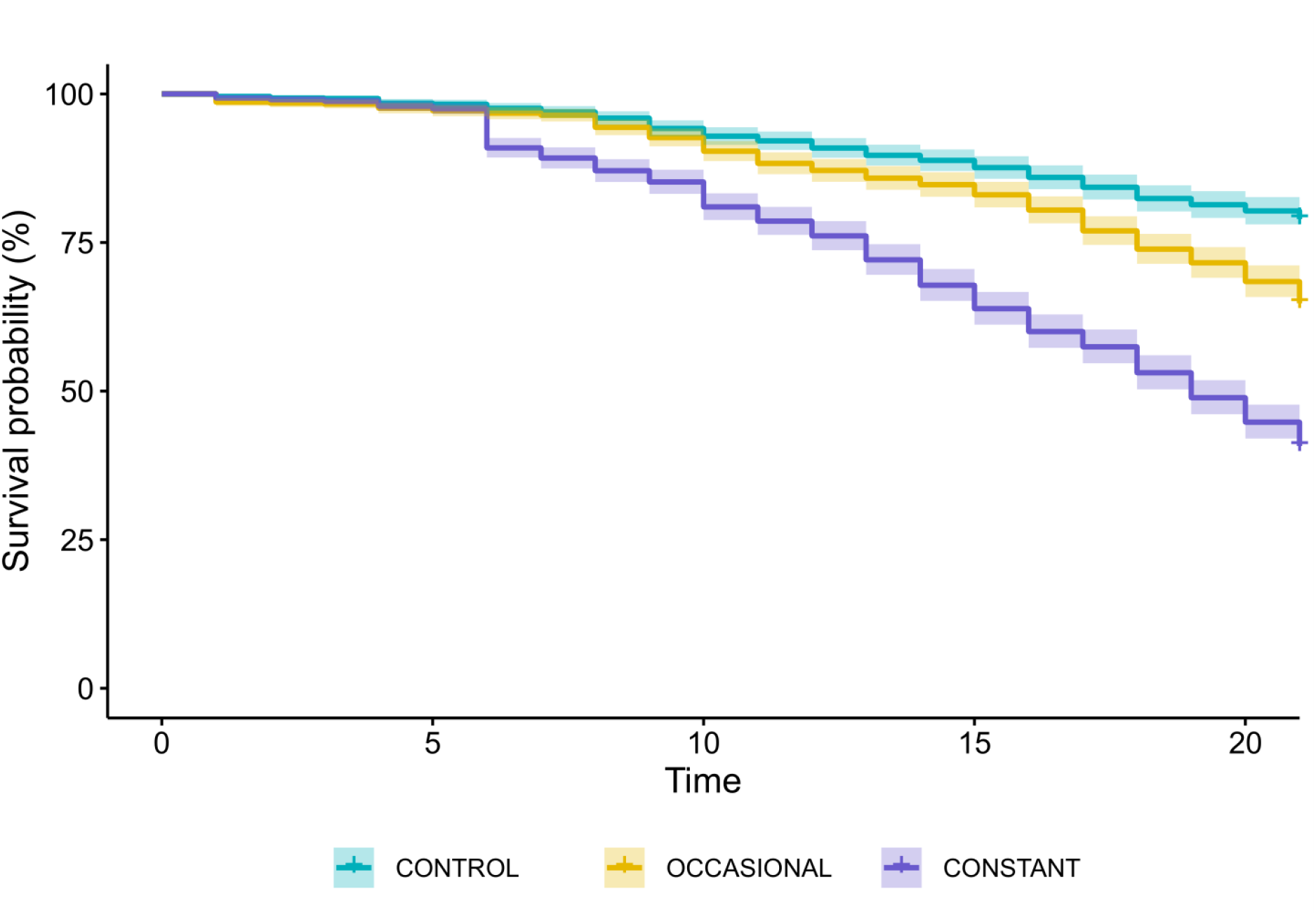
Survival plot for honey bees exposed to different types of diets for 21 days (CONTROL = the control groups exposed to pure 2 M sucrose solution, OCCASIONAL = the groups with occasional exposure to ethanol (addition of 1% ethanol in food every third day), CONSTANT = the groups with constant exposure to ethanol (food with the addition of 1% ethanol)). There were 12 cages for each group, each with an initial number of individuals ranging from 94-100. Shadings show 95% CI.

**Figure 3.**
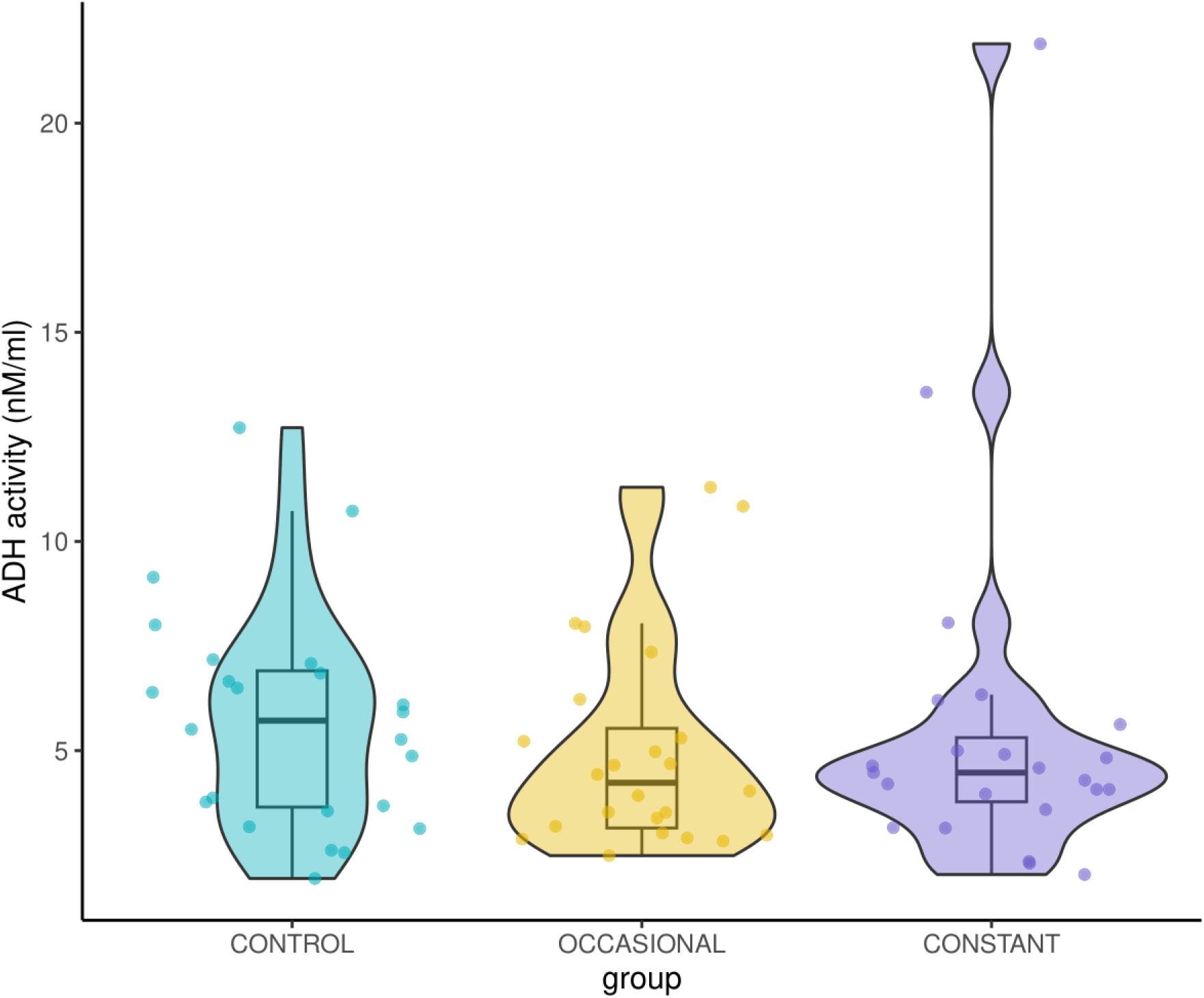
Violin plot for the ADH activity (mU / mL) in honey bees exposed to different types of diets for 21 days (CONTROL = the control groups exposed to pure 2 M sucrose solution (n = 24), OCCASIONAL = the groups with occasional exposure to ethanol (addition of 1% ethanol in food every third day) (n = 24), CONSTANT = the groups with constant exposure to ethanol (food with the addition of 1% ethanol) (n = 23)). Boxes indicate the median and interquartile range. Whiskers reach the smallest and largest values within 1.5 of the interquartile range. Coloured dots show individual data points.

## Discussion

Honey bees surely encounter and consume ethanol in their natural environment. They rely on ethanol for synthesising one of their crucial pheromones (Leoncini et al. 2004, Castillo et al. 2012a, Castillo et al. 2012b), possess enzymes to break down ethanol (Miler et al. 2021), and willingly consume it when mixed with sugars (Abramson et al. 2000, Abramson et al. 2004a, Maze et al. 2006, Sokolowski et al. 2012, Mustard et al. 2018). While previous research has focused on the effects of single and often high-concentration ethanol exposure on honey bees, this study investigates the consequences of repeated exposure to low levels of ethanol. Our findings reveal a significant decrease in lifespan due to ethanol exposure. Mortality rates in our study show that a significant drop in survival starts to appear after about 6 and 10 days of constant and occasional ethanol consumption, respectively. It is worth noting, however, that in natural conditions the average lifespan of forager-age bees is about a week (Dukas 2008). Therefore, while our results highlight the potentially harmful effects of even low ethanol concentrations and suggest cumulative impacts with each subsequent exposure, they need to be put in an ecological context. It seems likely that bees die for other reasons before the negative impact of routine, low-level ethanol exposure in their food appears. For comparison, in the study of Mustard et al. (2019), no discernible differences in mortality were observed in bees exposed daily to 1.25% ethanol and those abstinent. In their study, however, the authors used foragers of unknown age captured at hive entrances and a vast majority of all these bees died within a week. It is conceivable that in that study, the effects of diet had no chance to manifest (Mustard et al. 2019). Rasmussen et al. (2021) utilised overwintering bees with a longer-lived phenotype and demonstrated that 1% ethanol in their diet significantly decreased survival, but with a drop in mortality occurring only after a few days of ethanol exposure. In our study, even if we consider mortality over the entire 3 weeks of measurement, well over a third of all individuals survived daily ethanol consumption (Fig 2). Together, these results highlight the extraordinary resilience to the potentially toxic effect of ethanol in honey bees. Previous research has demonstrated that ethanol induces an increase in heat shock protein concentration in honey bee brain tissue (Hranitz et al. 2010). Rasmussen et al. (2021) also showed that ethanol intake disrupts their DNA methylation patterns. These effects likely constitute the basis of the eventual harmfulness of ethanol consumption and its accumulative toxicity. Yet, it takes time for such effects to reflect on the survivorship of bees.

Bees likely ingest ethanol with their food as they collect floral nectar. Another potential source of ethanol is presented in spoiled or overripe fruits (e.g. Dudley 2002, Dominy 2004, Sánchez et al. 2004) because bees often collect juices from such fruits, especially during periods without major flowering events. Our understanding of the sources of honey bee exposure to ethanol, as well as the frequency and concentration at which honey bees come into contact with this compound, remains a gap that requires extensive research. The same is true for other pollinating nectarivores as well as frugivores, all of which probably meet with ethanol routinely (Dudley and Maro 2021).

Our results revealed no differences in the ADH activity between the groups of individuals (Fig 3), suggesting that foragers consistently produce the enzyme, regardless of their ethanol exposure. This result is surprising, as it contradicts the expectation that prolonged alcohol consumption would induce ADH expression, as observed in vertebrates (Buris et al., 1985; Kishimoto et al., 1995; Tran et al., 2015). Notably, however, there are different types of ADH (Hernández-Tobías et al. 2011). In the case of fruit flies, in which ADH activity is closely related to alcohol tolerance and preferences (Ogueta et al. 2010), ADH type II was the focus of the study. Short-chain type II ADH is more typical for insects and might show higher activity with ethanol in bees, similar to *Drosophila* (Hernández-Tobías et al. 2011). In the case of honey bees, types I and II are present (Miler et al. 2021, Miler et al. 2022), but only the former is analysed here. Hence, our results regarding ADH activity cannot be treated as entirely conclusive. Similar results to ours were obtained in fruit-feeding butterflies, in which a diet containing ethanol did not alter ADH levels (Beaulieu et al. 2017). Miler et al. (2021) demonstrated ADH presence in honey bee foragers but not in intranidal workers that stay inside the hive and care for brood. Additionally, another study showed that even repeated ethanol exposure failed to induce ADH production in intranidal workers (Miler et al. 2022). Our current findings complement this prior research (Beaulieu et al. 2017, Miler et al. 2021, Miler et al. 2022) indicating that ADH type I activity remains constant, regardless of ethanol intake. The analysed type I ADH might not be restricted in its function to ethanol metabolism and this may confound the results. Additionally, considering our small sample size, we might have failed to detect an existing effect. We recommend future research to focus on this issue, including the activity of *Drosophila*-type ADH in bees.

Our findings stress the necessity for further research to find ecologically relevant levels of ethanol and the frequency of exposure to this substance in bees as well as their responses to this dietary component. The way for the comparative biology of ethanol exposure and response is open (Dudley and Maro 2021) and honey bees show promise in this context. Considering that ethanol is one of many neuroactive substances present in floral nectar that potentially mediate plant-pollinator interactions (Mustard 2020), this research field needs much more attention. It’s important to acknowledge that much of the existing research, including our own, has been conducted under laboratory conditions. Although such studies have limitations, laboratory cage trials in the study of the honey bee remain a commonly utilised research method, offering high comparability between groups and minimising the influence of external factors. However, this controlled environment may not fully capture the complexities of real-world scenarios. Therefore, to gain a more comprehensive understanding of the impact of alcohol on bees, it’s essential to complement laboratory studies with field research.

## Notes

Funding: This work was supported by the National Science Centre, Poland [grant number Sonata UMO-2021/43/D/NZ8/01044].

### Competing Interest Statement

The authors have declared no competing interest.

### Summary of Updates

Treatment group names were changed for clarity. Additional methodological details were provided. The introduction was updated to provide more background on the reason behind studying ADH. The experimental scheme was added (new Figure 1) and Figure 3 was revised to include additional information. Additional comments were included in the Discussion section.

